# Altered ultrasonic vocalizations in mouse models of FOXG1 syndrome

**DOI:** 10.64898/2026.11.25.769311

**Authors:** Sevda Abdavinejad, Dongjun Shin, Eunjin Kwak, Hohyun Hwang, Shin Jeon, Payton Maltbie, Soo-Kyung Lee, Micheal L. Dent, Valli Duvvuru

**Affiliations:** Department of Psychology, College of Arts and Sciences, University at Buffalo, Buffalo, New York, USA; Department of Biological Sciences, College of Arts and Sciences, University at Buffalo, Buffalo, New York, USA; FOXG1 Research Center, University at Buffalo, Buffalo, New York, USA; Department of Systems Pharmacology and Translational Therapeutics, University of Pennsylvania, Philadelphia, USA

**Keywords:** FOXG1, W300X, NEX-Cre, Ultrasonic Vocalizations, Autism Spectrum Disorder

## Abstract

FOXG1 syndrome is a severe neurodevelopmental disorder with communication impairment as a defining clinical feature, yet vocal behavior has not been characterized in animal models of FOXG1. We examined neonatal ultrasonic vocalizations in two mouse models that disrupt *Foxg1* in distinct developmental and cellular contexts: a NEX-Cre-driven conditional heterozygous model targeting postmitotic cortical and hippocampal excitatory neurons, and a W300X/+ knock-in model carrying a patient derived truncating mutation at the endogenous *Foxg1* locus. Vocalizations were recorded at postnatal day 7 and analyzed for call number, duration, frequency range, and repertoire composition. The two models exhibited distinct vocal phenotypes. Conditional heterozygous mice produced longer calls while total vocal output, frequency range, and repertoire composition were preserved. In contrast, W300X/+ mice produced shorter calls, showed a significant shift in repertoire composition, and a non-significant trend toward reduced vocal output. The effect of genotype on call duration differed significantly between models, demonstrating opposing effects on temporal vocal structure. Frequency range was not significantly altered by genotype in either model, although sex was associated with frequency range in the conditional heterozygous cohort. These findings provide the first evidence that *Foxg1* disruption alters neonatal vocal behavior and show that the resulting phenotype depends on the developmental, cellular, and molecular context of the mutation. Neonatal vocalizations therefore provide a sensitive behavioral readout for distinguishing functional consequences of disease relevant FOXG1 perturbations.

## Introduction

### FOXG1 syndrome

Forkhead Box G1 (FOXG1, previously known as Brain Factor-1) is a winged-helix transcription factor with indispensable roles in telencephalic development [1]. Homozygous loss of *Foxg1* is embryonically lethal in mice [2,3], while *de novo* heterozygous pathogenic variants cause FOXG1 syndrome (FS), a rare neurodevelopmental disorder, in humans. FS is characterized by cortical malformations, including corpus callosum hypogenesis, delayed myelination, hippocampal malformation, simplified gyral patterns, and microcephaly [4–6]. Clinically, FS presents with epilepsy, severe intellectual disability, sleep disturbances, and irritability [7–9]. Among the most prominent features of the syndrome is a profound deficit in communication: the majority of individuals with FS do not develop spoken language, with only approximately 20% acquiring any spoken words, and most remaining entirely nonverbal throughout their lives [7,8]. Individuals with FS also commonly exhibit features associated with autism spectrum disorder (ASD), including impaired social interactions, stereotypic movements, and difficulty with eye contact [5]. Despite the centrality of communication deficits to the clinical presentation of FOXG1 syndrome, the impact of FOXG1 disruption on vocal communication has never been examined in any animal model.

### Rodent models of FOXG1 syndrome

Extensive research using mouse models has established essential roles for FOXG1 throughout telencephalic development. FOXG1 regulates cortical neurogenesis by controlling progenitor proliferation and differentiation within the subventricular and ventricular zones [10–12]. It also guides neuronal migration to form cortical layers and plays a crucial role in axonal projection formation and maintaining the balance of excitatory and inhibitory circuits [13,14]. More recent patient-specific FOXG1 models have extended these findings by showing that disease-associated variants can disrupt radial glial identity, neuronal migration, axonal organization, synaptic gene programs, and postnatal myelination [6].

Conditional models have been particularly useful for defining the postmitotic functions of FOXG1 within cortical and hippocampal excitatory neurons. NEX-Cre-mediated deletion of *Foxg1* disrupts cortical and hippocampal development and long-range connectivity, with heterozygous loss producing milder but still detectable structural abnormalities compared with conditional knockout models [13]. By contrast, the W300X/+ knock-in model carries a patient-derived truncating mutation at the endogenous *Foxg1* locus, allowing FOXG1 dysfunction to arise across its broader developmental expression domain [15]. Together, these models provide complementary approaches for examining how restricted postmitotic haploinsufficiency and a disease-associated truncating mutation influence early behavioral phenotypes.

### Rodents as models for communication disorders

Mice produce ultrasonic vocalizations (USVs) in a number of contexts for communication. USVs provide a sensitive behavioral readout for investigating the developmental and genetic mechanisms underlying social and communicative behaviors [16–19]. Neonatal mice emit USVs when separated from the dam or experience temperature stress, with calling rates reaching their peaks between P7–9 [20–23]. Both the rate and acoustic structure of pup USVs are influenced by genetic background and developmental stage, making them useful for detecting early alterations in vocal behavior.

USV abnormalities have been reported across multiple mouse models of neurodevelopmental disorders associated with human communication and language impairments, including *Foxp2* [24–26], *Shank* family [27–29], and *Cntnap2* [30] models, supporting the use of USVs as quantitative measures of early behavioral phenotypes. USV production and modulation depend on distributed neural circuits, including the periaqueductal gray and forebrain regions such as medial prefrontal, motor, and cingulate cortices [31,32], making neonatal USVs particularly relevant for detecting early circuit-level abnormalities before mature behavioral phenotypes are established. In neonatal mice, brain-wide activity mapping during maternal-separation USVs implicates a distributed network spanning cortical, limbic, and brainstem regions [33]. Cortico-striatal circuitry has also been implicated in neonatal vocalizations [34,35].

Together, these findings indicate that early vocal behavior depends on coordinated activity across multiple forebrain and subcortical circuits. Given the broad role of FOXG1 in telencephalic development, disruption of these networks provides a plausible basis for altered vocal behavior in FOXG1 syndrome models.

To investigate how *Foxg1* disruption affects early vocal behavior, we characterized USVs in two FOXG1 syndrome-associated mouse models at P7, within the developmental period of peak USV emission in neonatal mice [36,37], in a NEX-Cre-driven conditional heterozygous model and the W300X/+ knock-in model. By examining these models in parallel, we sought to determine whether distinct forms of *Foxg1* disruption produce shared or model-specific alterations in neonatal vocalizations.

## Materials and methods

### Ethics statement

All procedures were approved by the University at Buffalo, SUNY’s Institutional Animal Care and Use Committee (IACUC) under protocol 202000055.

### Experimental model and subject details

Male and female mice from two *Foxg1*-related genetic models were used in the study. The *Foxg1*cHet cohort included 25 mice: *Foxg1*cHet mice (n = 5 F, 6 M) and *Foxg1*^f/+^ controls (n = 9 F, 5 M). The W300X cohort included 33 mice: W300X mice (n = 6 F, 4 M) and C57BL/6 wild-type controls (n = 15 F, 8 M).

The mice were housed at a 12 h dark/12 h light cycle, 25°C, and 50% humidity. All housing and analyses complied with the guidelines of the IACUC of the University at Buffalo. Foxg1 flox [13] mice and Nex-Cre mice [38] have been described previously. All experiments were conducted on mice with the C57BL/6 background, and Foxg1-cHet mice were generated by crossing Foxg1flox/flox with Nex-Cre/+ mice. Foxg1 W300X/+ knock-in mouse line, carrying the mouse equivalent of the human FOXG1 W308X pathogenic variant was generated and characterized as previously described [15]. Foxg1 W300X/+ mice were crossed with C57BL/6 wild-type mice to generate W300X/+ heterozygous offspring and wild-type littermate controls.

The pups used in this study were part of a larger neonatal ICV cohort and therefore received PBS control injections within 36 h after birth. Mutant and control pups within each cohort received identical injections, so the procedure was balanced across genotypes. ICV injection was performed under cryoanesthesia and using 10 μL Hamilton micro-syringe into the lateral ventricle of a single hemisphere. The pups remained with the mother from birth till weaning and underwent USV recordings on P7.

### Recording apparatus

The recording chambers were standard mouse cages (30 x 19 x 13 cm) lined with fresh corn cob bedding. These cages were placed at the center of a double-walled sound-attenuated booth lined with anechoic foam (10.2 cm Sonex, Illbruck Inc., Minneapolis, MN). Ultrasonic condenser microphones (Avisoft Bioacoustics, model CM16/CMPA, Berlin, Germany) were placed above the recording cage, pointing downwards, about 50 cm from the bottom of the cage. The microphones have a flat frequency response (± 6 dB) between 25 and 150 kHz. Recordings were made with Avisoft UltraSoundGate 116H recorders at a 300 kHz sampling rate and 16-bit resolution, connected to a Hewlett Packard (HP) notebook running Avisoft-RECORDER software.

### USV recording and analysis

P7 mice were left to acclimatize to the recording room for 1 hour prior to the experiment. At the time of the experiment, each individual mouse was placed in the recording cage without the lid and allowed to habituate for 30 seconds. This was followed by a 1-minute-long recording period, following which the mouse’s weight was recorded. The recording files were then exported in .wav format. USV identification and categorization were performed manually by expert raters (SA and VD) using Raven Lite (v. 2.0, Cornell Lab of Ornithology) [39]. Recordings from the two cohorts were acquired and scored in parallel over the same period by the same two raters, so cohort and genotype were not confounded. Inter-rater reliability for total call counts per recording was assessed using a two-way mixed-effects intraclass correlation coefficient (ICC) with absolute agreement (ICC(A,1) = 0.998, 95% CI [0.994, 0.999], *p* < 0.001). Calls were categorized into spectrotemporal types based on previously published criteria: chevron, chirp, complex, downsweep, flat, harmonic, inverse chevron, jump, and upsweep [40], plus an unstructured call category defining calls that could not be categorized into the other nine clear categories [41].

### Statistical Analysis

All analyses were performed in R version 4.5.1 (R Core Team, 2025), with the individual mouse treated as the experimental unit. Tests were two-sided with significance set at *p* < 0.05. All mice were included in total-call analyses; acoustic and repertoire analyses included only vocalizing mice (*Foxg1*cHet: mutants, n = 5 F, 6 M; controls, n = 9 F, 5 M; W300X: mutants, n = 5 F, 2 M; WT controls, n = 13 F, 8 M). Three of 10 W300X mutants and 2 of 23 W300X controls produced no calls, so the acoustic and repertoire results for that cohort are conditioned on vocalizing animals.

Total USV counts were analyzed separately within each model using negative binomial generalized linear models with genotype, sex, and their interaction. Genotype-by-sex interactions were tested first; when not significant, additive models were used for inference. Genotype and sex effects were assessed by likelihood-ratio testing, and estimated marginal means and planned genotype contrasts were obtained from the fitted models. Acoustic analyses were performed on mouse-level median call duration and median frequency range using linear models with genotype and sex. Interactions were tested first, followed by additive models when not significant. HC3 heteroskedasticity-robust covariance estimates were used for W300X acoustic analyses and for cross-model analyses. Cross-model models included genotype status, model, sex, and the genotype-status-by-model interaction.

Vocal repertoire was analyzed from mouse-level relative abundances of 10 call categories using Bray-Curtis dissimilarities. Genotype, sex, and genotype-by-sex effects were tested by PERMANOVA with 9,999 permutations, followed by additive models when the interaction was not significant. Multivariate dispersion was assessed with PERMDISP, and Lingoes correction was applied for dispersion analyses and principal coordinates analysis (PCoA). PCoA was used for visualization only.

## Results

### Total USV output

We first examined whether *Foxg1*-related mutations altered overall USV production. In *Foxg1*cHet mice, total call production did not differ between Control and mutant animals (negative-binomial model, genotype effect, *p* = 0.734; Fig. 1A). Model-estimated mean call counts were 80.9 calls in Controls and 72.8 calls in mutants. Sex was also not significantly associated with vocal output (*p* = 0.848).

**Fig 1.**
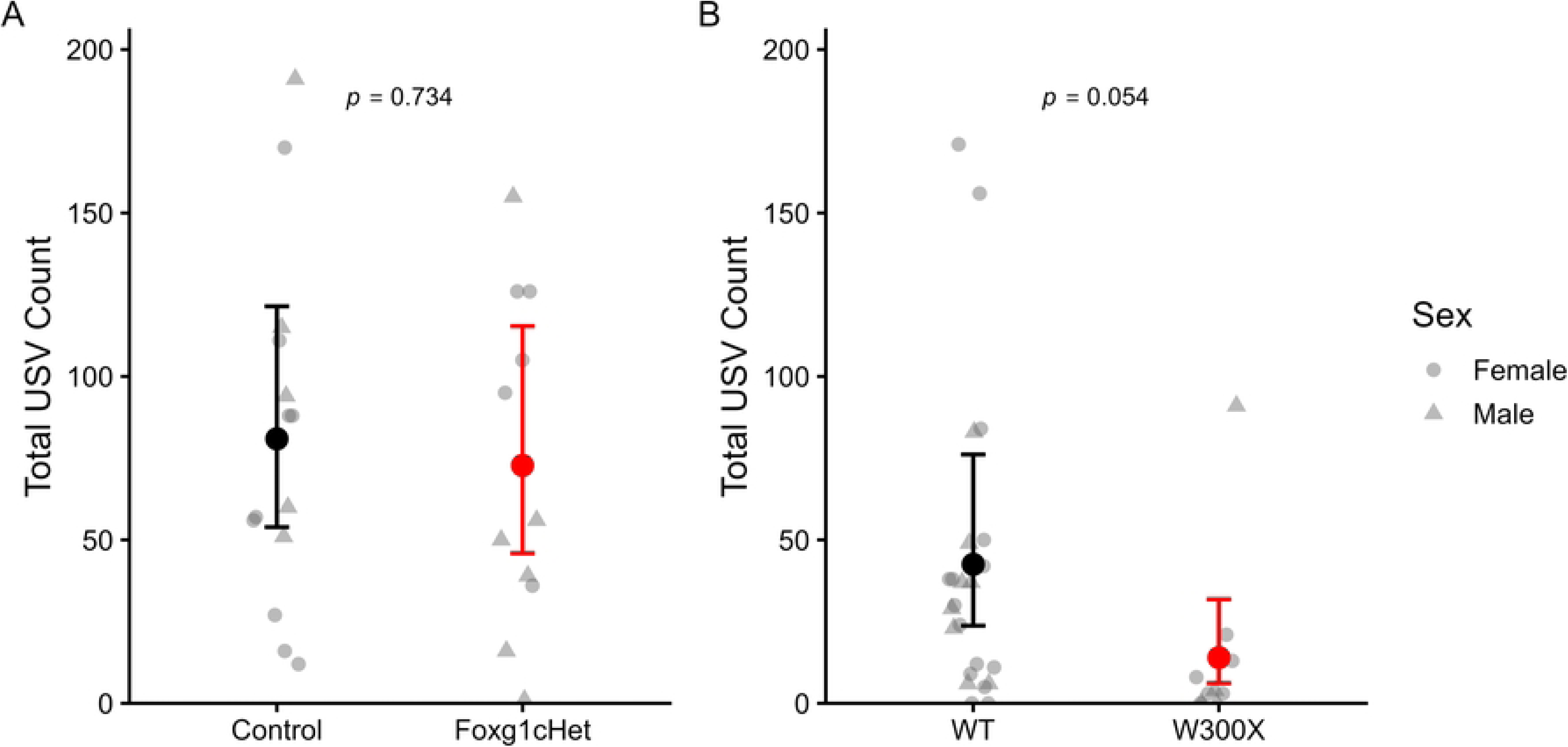
Total USV output in *Foxg1*cHet and W300X mice. (A) Total number of USVs emitted by *Foxg1*cHet mice and *Foxg1*^f/+^ controls during the recording period. No significant genotype effect was detected. (B) Total USV number in W300X mice and WT controls. Vocal output was lower in W300X mice than in controls, although the overall genotype effect did not reach statistical significance. Small points represent individual mice. Large symbols represent model-estimated marginal means, and error bars indicate 95% confidence intervals.

W300X mice showed lower estimated vocal output than WT controls (Fig. 1B). The omnibus likelihood-ratio test for genotype did not reach conventional statistical significance (*p* = 0.054). The corresponding model-estimated Control-versus-W300X contrast was nominally significant (*p* = 0.035), but these two tests address the same hypothesis, so we interpret this difference as a trend toward reduced vocal output rather than a significant genotype effect. Estimated mean call counts were 42.5 in Controls and 14.0 in mutants. This corresponded to an estimated Control-to-W300X call-count ratio of approximately 3.04. The mutant estimate is based on 10 animals, three of which produced no calls and one of which produced 91, more than four times any other mutant; excluding that animal, the genotype effect is significant (*p* = 0.00063). Sex was not significantly associated with total vocal output in the W300X cohort (*p* = 0.728).

### Call duration

We next examined mouse-level median call duration. *Foxg1*cHet mice produced longer calls than Controls (genotype effect, *p* = 0.032; Fig. 2A), with model-estimated durations of 34.2 ms in Control mice and 47.2 ms in *Foxg1*cHet mice. A significant effect of sex was also detected (*p* = 0.024), whereas the genotype-by-sex interaction was not significant (*p* = 0.862).

**Fig 2.**
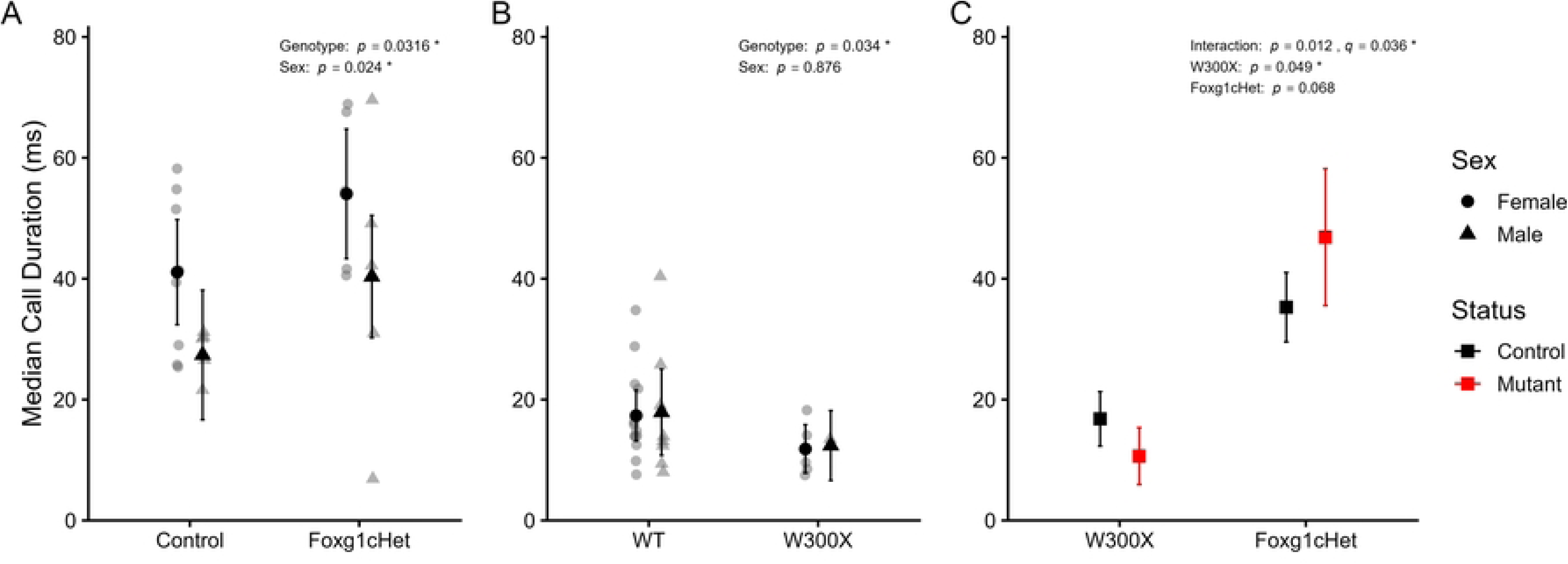
Call duration differs between Foxg1cHet and W300X mice. (A) Median USV duration in *Foxg1*cHet mice and *Foxg1*^f/+^ controls. *Foxg1*cHet mice produced longer calls than controls, and males produced shorter calls than females. The genotype-by-sex interaction was not significant. (B) Median USV duration in W300X mice and C57 WT controls. W300X mice produced shorter calls than controls, with no significant sex effect or genotype-by-sex interaction. (C) Cross-model comparison of genotype effects on call duration. The effect of genotype differed significantly between models. Small points in panels A and B represent individual mice. Large symbols and error bars in panels A and B represent model-estimated marginal means and 95% confidence intervals. Panel C shows model-estimated marginal means, averaged over sex, with 95% confidence intervals.

The direction of the genotype effect was reversed in W300X mice (Fig. 2B). W300X mice produced shorter calls than Controls, with model-estimated durations of 12.1 ms in W300X mice and 17.6 ms in Controls. The genotype effect was significant using HC3 heteroskedasticity-robust inference (*p* = 0.034), whereas sex was not significant (*p* = 0.876). There was also no evidence of a genotype-by-sex interaction (*p* = 0.889).

Consistent with these opposing within-model patterns, the combined analysis revealed a significant genotype-status-by-model interaction (*p* = 0.012; Fig. 2C), indicating that the relationship between genotype status and call duration differed across the two models. Within the combined model, the estimated Control-versus-mutant contrast was significant for W300X (*p* = 0.049) but did not reach significance for *Foxg1*cHet (*p* = 0.068).

### Frequency range

We next analyzed mouse-level median frequency range. In *Foxg1*cHet mice, frequency range did not significantly differ between Control and mutant animals (genotype effect, *p* = 0.567; Fig. 3A). In contrast, a significant effect of sex was observed (*p* = 0.004), with females showing broader frequency ranges than males. The genotype-by-sex interaction was not significant (*p* = 0.598). In the W300X cohort, neither genotype nor sex significantly affected frequency range (genotype, *p* = 0.296; sex, *p* = 0.649; Fig. 3B).

**Fig 3.**
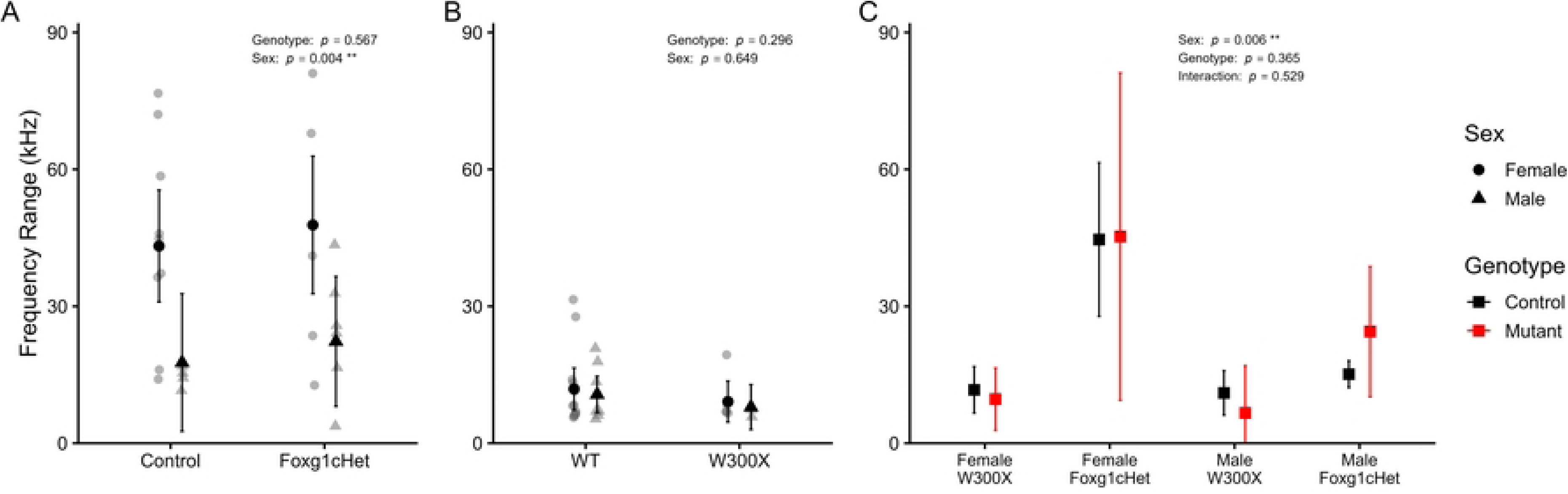
Median frequency range in Foxg1cHet and W300X mice. (A) Median USV frequency range in *Foxg1*cHet mice and *Foxg1*^f/+^ controls. No significant genotype effect was detected, whereas a significant sex effect was observed, with females exhibiting broader frequency ranges than males. The genotype-by-sex interaction was not significant. (B) Median USV frequency range in W300X mice and C57 wild-type controls. Neither genotype nor sex significantly affected frequency range, and the genotype-by-sex interaction was not significant. (C) Cross-model comparison of frequency range. No significant Status × Model interaction was detected. A significant overall sex effect was present across models. Frequency range is expressed in kHz. Small points in panels A and B represent individual mice. Large symbols and error bars in panels A and B represent model-estimated marginal means and 95% confidence intervals. Panel C shows observed subgroup means with 95% confidence intervals.

In the combined cross-model analysis, sex remained significantly associated with frequency range (*p* = 0.006; Fig. 3C). There was no significant overall effect of genotype status (*p* = 0.365) and no genotype-status-by-model interaction (*p* = 0.529). Overall, frequency range showed no genotype-associated shift in either model, whereas sex was significantly associated with this measure.

### Vocal repertoire composition

We next examined whether genotypes altered the relative distribution of vocalization types. For each mouse, categorized calls were converted to relative proportions across ten call types, and overall repertoire dissimilarity was quantified using Bray-Curtis distances.

In *Foxg1*cHet mice, overall repertoire composition did not significantly differ between genotypes (PERMANOVA: *F1*,22 = 0.781, *R^2^* = 0.033, *p* = 0.575; Fig. 4A, B). There was also no significant effect of sex (*F1*,22 = 1.042, *R^2^* = 0.043, *p* = 0.381) and no genotype-by-sex interaction (*F1*,21 = 0.827, *R^2^* = 0.035, *p* = 0.539). Multivariate dispersion did not differ among genotype-by-sex groups (Lingoes-corrected PERMDISP, *p* ≈ 0.291), indicating no evidence that unequal within-group dispersion confounded the PERMANOVA results.

**Fig 4.**
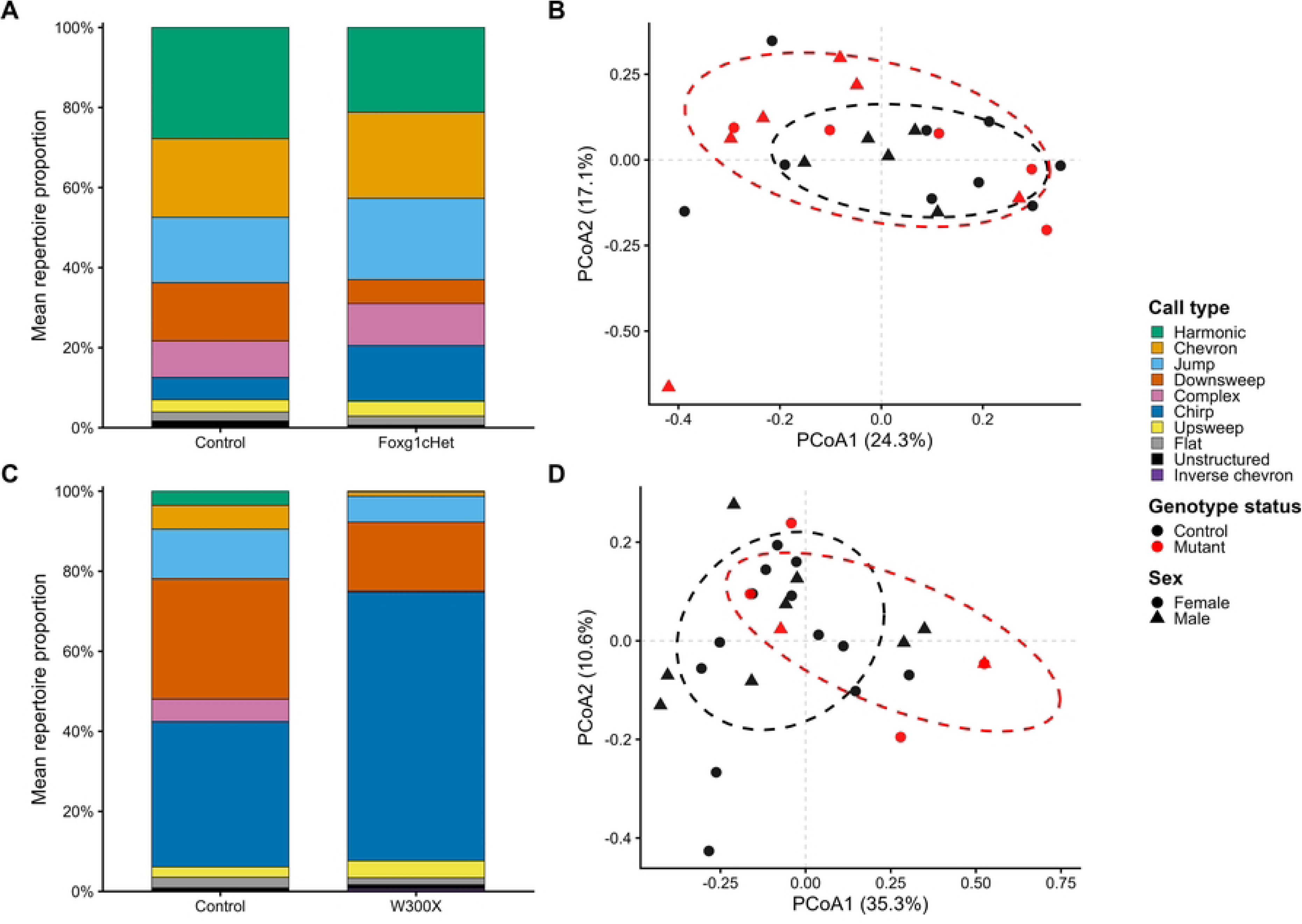
Vocal repertoire composition differs between *Foxg1*cHet and W300X mice. (A) Relative distribution of USV call categories in *Foxg1*cHet mice and *Foxg1*^f/+^ controls. No significant effects of genotype, sex, or the genotype-by-sex interaction on repertoire composition were detected. (B) Principal coordinates analysis (PCoA) of Foxg1cHet repertoire composition based on Lingoes-corrected Bray–Curtis distances; PCoA1 and PCoA2 explained 24.3% and 17.1% of the positive eigenvalue variance, respectively. (C) Relative distribution of USV call categories in W300X mice and C57 WT controls. Repertoire composition differed significantly by genotype (*p* = 0.013), whereas no significant sex or genotype-by-sex interaction was detected. (D) PCoA of W300X repertoire composition based on Lingoes-corrected Bray–Curtis distances; PCoA1 and PCoA2 explained 35.3% and 10.6% of the positive eigenvalue variance, respectively.

In contrast, W300X mice showed a significant genotype-associated shift in overall repertoire composition relative to Controls (PERMANOVA: *F1*,25 = 4.299, *R^2^* = 0.146, *p* = 0.013; Fig. 4C,D). Genotype therefore accounted for approximately 14.6% of the variation in Bray-Curtis repertoire dissimilarity. Sex was not significantly associated with repertoire composition (*F1*,25 = 0.176, *R^2^* = 0.006, *p* = 0.937), and there was no genotype-by-sex interaction (*F1*,24 = 0.033, *R^2^* = 0.001, *p* = 0.990). Multivariate dispersion was also comparable among genotype-by-sex groups (Lingoes-corrected PERMDISP, *p* = 0.852).

## Discussion

Communication deficits are among the most prominent clinical features of FOXG1 syndrome, with the majority of affected individuals lacking spoken language entirely [7,8]. Despite this, no prior study has examined vocalization in any *Foxg1* mouse model. The present study addresses this gap by providing the first characterization of USVs in two FOXG1 syndrome models: a conditional heterozygous model and the W300X/+ knock-in model that mimics the W308X/+ human mutation seen in FS [15].

*Foxg1*cHet mice showed increased call duration while total vocal output, median frequency range, and repertoire composition were preserved. In contrast, W300X/+ mice exhibited shorter calls, altered repertoire composition, and a trend toward reduced vocal output. Frequency range was not significantly altered by genotype in either model, although a significant sex effect was observed in the *Foxg1*cHet cohort. Together, these findings indicate that FOXG1 disruption does not produce a uniform vocal phenotype but instead affects distinct dimensions of neonatal vocal behavior depending on the genetic and developmental context of the perturbation.

USV abnormalities have been reported across other neurodevelopmental disorder models with non-uniformly affected features [17,42]. *Shank* models show varying quantitative and qualitative abnormalities, including sex-dependent genotype effects [43], while BTBR pups exhibit increased call production and altered vocal characteristics relative to C57BL/6J mice [44]. USV structure is also influenced by developmental, genetic, behavioral, and social context [40,45,46], including modulation of isolation-induced vocalizations by familiar social odors [47]. Individual USV dimensions can likewise be affected independently: Dock4 knockout pups show altered developmental repertoire organization [48], valproate-exposed and Caps2−/− pups show shortened call duration whereas Chd8+/− mice exhibit comparatively limited acoustic alterations [49], and disruption of Foxp2 reduces pup vocalization while leaving call duration, peak frequency, and bandwidth relatively preserved [24]. Together, this literature provides context for the distinct phenotypes observed here and supports interpreting the opposing effects on call duration in Foxg1cHet and W300X/+ mice as qualitatively different rather than as different degrees of a common impairment.

Sex also contributed to the Foxg1cHet phenotype, with females exhibiting broader frequency ranges than males despite no overall genotype effect on this measure. Sex-dependent variation in neonatal USVs changes across development; a recent meta-analysis identified developmental stage as a significant moderator of sex differences in call duration, with control females producing longer calls than males during early postnatal development and the direction reversing later [50]. These findings reinforce the importance of considering sex and developmental stage when interpreting neonatal USV phenotypes.

We generated Foxg1 conditional heterozygous mice for our study using a *NEX-Cre* driver, which restricts *Foxg1* deletion to postmitotic cortical and hippocampal neurons, sparing other *Foxg1-*expressing populations including inhibitory interneurons, neural progenitors, and subcortical regions. The cHet model showed increased call duration but no detectable genotype effect on total call number, median frequency range, or repertoire composition. Such selective effects are consistent with previous genetic models in which individual components of mouse USVs are differentially affected rather than changing in parallel [51–53]. These findings demonstrate that *Foxg1* dysfunction restricted to postmitotic excitatory neurons alone is sufficient to alter neonatal vocal behavior, implicating the integrity of cortical and hippocampal circuitry in the temporal structure of neonatal vocalizations.

Whereas the cHet model isolates Foxg1 dysfunction to a defined postmitotic neuronal population, the W300X mutation is present throughout the endogenous Foxg1 expression domain [54] and therefore acts from the earliest stages of telencephalic development [15]. FOXG1 plays established roles in neural progenitor proliferation [11,12], neuronal specification [10,55,56], migration [55], and telencephalic morphogenesis and regional patterning [2,57,58]. In addition, FOXG1 is also expressed in subcortical telencephalic populations, including the striatum [59]. This raises the possibility that W300X affects developmental processes and neural populations that are not targeted by the conditional manipulation. Truncating FOXG1 variants may additionally produce allele-specific molecular consequences that are not fully recapitulated by heterozygous loss alone, including loss of specific functional domains, and, as shown for FOXG1 Q84Pfs/+, interactions between truncated and full-length FOXG1 that alter its subcellular distribution [6]. In the W300X mouse model, the truncated FOXG1 protein has been reported to be more abundant and to show greater nuclear localization than the full-length protein, suggesting W300X pathogenicity may involve effects of the truncated protein in addition to the reduced FOXG1 dosage [15]. The broader W300X/+ vocal phenotype may therefore reflect this distinct developmental and allele-specific context.

Several limitations should be noted. The two models were compared against different control genotypes, *Foxg1*f/+ littermates for the cHet model and C57BL/6 wild-type littermates for W300X, and absolute call counts and durations differed between the two control groups; the within-model genotype comparisons are therefore the primary analyses, and the cross-model interaction should be read with this difference in baseline in mind. All pups in both cohorts received a neonatal PBS ICV injection as part of a larger experiment, so absolute USV values are not directly comparable to published cohorts of uninjected P7 mice, although the procedure was balanced across genotypes within each cohort. Recordings were obtained at a single postnatal age, which cannot distinguish a persistent deficit from a shift in developmental timing. Finally, calls were classified manually; although inter-rater agreement for total call counts was high, category assignment remains more subjective than automated classification.

## Conclusions

Distinct USV phenotypes observed across the cHet and W300X/+ models suggest that the behavioral consequences of *Foxg1* disruption are shaped by the developmental, cellular, and molecular context of the perturbation. This is particularly evident in the opposing effects on call duration, suggesting that the two models capture complementary aspects of FOXG1 disruption, with the cHet model isolating the consequences of cell-type restricted haploinsufficiency and the W300X model capturing broader developmental and allele-specific context. This distinction may be particularly relevant given evidence that neonatal vocalizations engage cortico-striatal circuitry [34,35], raising the possibility that disruption of both cortical and striatal components could contribute to the W300X phenotype. More broadly, the divergence between these models underscores the importance of considering model-specific developmental and molecular context when interpreting behavioral phenotypes in neurodevelopmental disorders.

## Data availability

All data underlying the findings described in this manuscript are provided in S1 Data.

## Author contributions

Sevda Abdavinejad: Conceptualization, Data Curation, Formal Analysis, Investigation, Methodology, Project Administration, Resources, Software, Supervision, Validation, Visualization, Writing – original draft, Writing – review & editing

Dongjun Shin: Conceptualization, Data Curation, Investigation

Eunjin Kwak: Data Curation, Investigation

Hohyun Hwang: Data Curation, Investigation

Shin Jeon: Visualization, Investigation

Payton Maltbie: Conceptualization, Data Curation, Investigation

Soo-Kyung Lee: Conceptualization, Funding Acquisition, Methodology, Project Administration, Resources, Supervision, Validation, Writing – review & editing

Micheal Dent: Conceptualization, Methodology, Project Administration, Resources, Supervision, Validation, Writing – review & editing

Valli Duvvuru: Conceptualization, Data Curation, Formal Analysis, Investigation, Methodology, Project Administration, Resources, Software, Supervision, Validation, Visualization, Writing – original draft, Writing – review & editing

## Supporting information

S1 Data. Per-animal USV measures. Total call counts, median call duration, median frequency range, and counts for each of the ten call categories for every mouse in both cohorts, with genotype and sex.

